# Rapid *in vitro* platform for functional analysis of maternal effect genes during mouse oocyte growth

**DOI:** 10.64898/2026.03.24.709698

**Authors:** Keisuke Sasaki, Yuhkoh Satouh, Mamoru Michizaki, Atsushi Jinno-Oue, Toshiyuki Matsuzaki

**Author notes:** **Correspondence**: Keisuke Sasaki, Ph.D. Faculty of Bioresources and Environmental Science, Ishikawa Prefectural University, 1-308 Suematsu, Nonoichi, Ishikawa 921-8836, Japan,.

## Abstract

Understanding the functions of maternal effect genes during oocyte growth is essential for elucidating the mechanisms of oogenesis and early embryonic development. However, conventional gene knockout and conditional knockout approaches require extensive breeding and are time-consuming. Here, we present a rapid *in vitro* gene functional analysis system that combines microinjection of mRNA, siRNA and plasmid DNA into mouse secondary follicles with a two-step oocyte growth culture system. Mouse secondary follicles were subjected to microinjection of mCherry mRNA and subsequently cultured for 15 days to produce fully grown oocytes. mCherry fluorescence persisted throughout the oocyte growth period but declined rapidly after fertilization. Despite minor cellular damage occasionally caused by microinjection, injected follicles developed normally and retained developmental competence. To evaluate the efficiency of gene suppression, we introduced siRNA targeting *Dnmt3l*, which is abundantly expressed during oocyte growth phase. Although *Dnmt3l* deficiency is known not to affect oocyte growth, we observed that oocyte growth was maintained normally despite a marked reduction in endogenous *Dnmt3l* mRNA levels in our knockdown model. These results demonstrate that this method enables efficient manipulation of gene expression specifically during oocyte growth while preserving developmental competence, providing a versatile platform for rapid functional screening of maternal effect genes *in vitro*.

## Introduction

In mammals, dormant oocytes in primordial follicles are maintained throughout reproductive age and parts of them are activated and grown to ovulate in response to endogenous sex hormone. During the growth phase, oocytes undergo various events including follicle growth, acquirement of epigenetic modification, and maternal factor accumulation [1–5]. In mice, gene knockout (KO) and germline-specific conditional knockout (CKO) strategies are commonly used to study gene functions involved in these processes [6–8]. However, there has been an issue that generation and maintenance of KO/CKO mice is time-consuming and needed multi-generational breedings.

To address this, we previously developed a novel oocyte-specific conditional gene knockdown (KD) system [9]. The CRISPR-mediated targeted insertion of artificial microRNA sequences into the intronic region of zona pellucida glycoprotein 3 (*Zp3*) gene enabled the generation of transgenic mice with an oocyte-specific KD effect, allowing us to analyze the functions of maternal effect genes more efficiently than with the traditional methods. Exogenous microRNA precursors are driven under control of *Zp3* transcription and processed into mature forms in oocytes entering to the growth phase, where they silence target genes complementary to artificial microRNA sequences.

In addition to the *in vivo* model, we also developed an *in vitro* approach involving low-invasive mRNA transfer to fully grown oocytes at the germinal vesicle stage [10]. Enclosed cumulus-oocyte complexes (COCs) were electroporated with mRNA and/or small interfering RNA (siRNA), then matured and fertilized, retaining developmental potential to the blastocyst stage. To disrupt maternal effect genes *in vitro*, it is necessary to suppress gene expression at earlier stages of oocyte growth, since siRNA introduction to fully grown oocytes are not able to degrade accumulated maternally translated proteins. Another *in vivo* electroporation method applied to whole ovaries does not deliver exogenous DNA into the growing oocytes but rather targets the oocyte-surrounding granulosa cells [11].

Other research groups also reported that microinjection of mRNA and/or siRNA directly into immature growing oocytes in secondary follicles and further simple follicle culture facilitated the understanding of maternal effect gene functions during oocyte growth [12–14]. However, it has been unclear whether their follicle culture systems could produce fertile oocytes developing to term. We previously established a robust *in vitro* oocyte growth (IVG) culture system using growing oocytes in the secondary follicles to produce fertile oocytes. [15–17]. In this study, we demonstrate a novel *in vitro* gene screening method which is consisted of mRNA and siRNA microinjection into the mouse secondary follicles and IVG. This technique facilitates time-efficient functional analysis of oocyte-associated genes while preserving developmental competence, all within a 16-day culture period.

## Results and discussion

We first designed a system to generate exogenous DNA or mRNA-delivered fully grown oocytes by combining microinjection with a two-step IVG (**Fig. 1A–B**). Following the previously reported method [12], secondary follicles isolated from female mice at postnatal day 10 (P10) were cultured in petri dishes for 1 day. Newly isolated secondary were often surrounded by adherent stromal cells, making them technically challenging to manipulate by microinjection; however, during this initial culture, these stromal cells adhered to the culture dish and were effectively removed from the follicles, which facilitated subsequent microinjection. To trace the expression of the injected nucleic acids in the growing oocytes, fluorescent reporters were utilized. We microinjected either a plasmid DNA construct expressing mCherry under the control of the *Stella* promoter into the nucleus of growing oocytes, or mCherry mRNA into the cytoplasm of the growing oocytes. For both delivery methods, robust fluorescence was observed 2 days post microinjection (**Fig. 1C**). Given that nuclear microinjection is technically demanding and physically risks compromising nuclear integrity, we concluded that cytoplasmic mRNA microinjection is more straightforward and less invasive than the nuclear injection. Consequently, we adopt cytoplasmic microinjection for all subsequent experiments.

**Figure 1.**
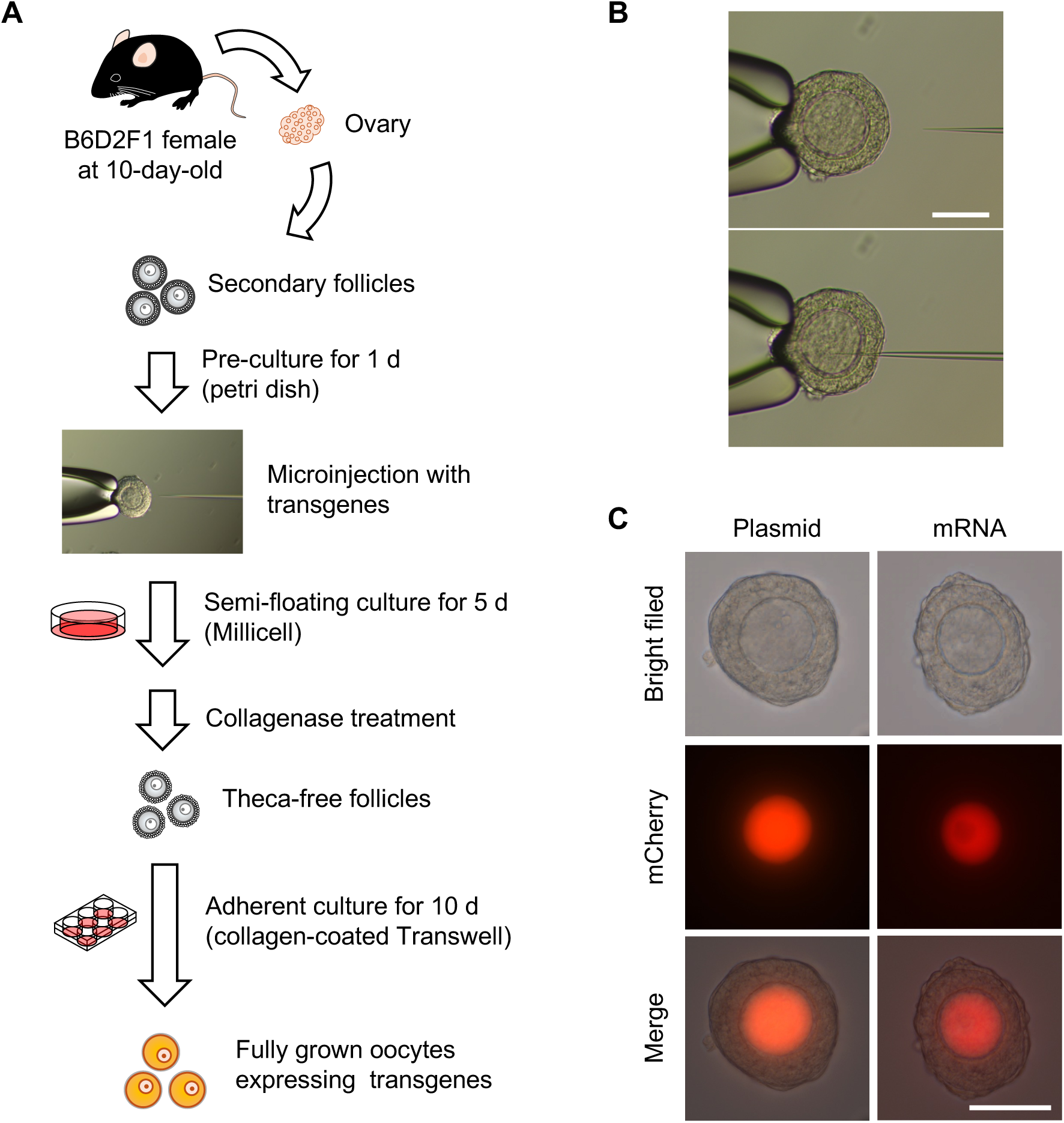
Experimental design for microinjection and *in vitro* growth of mouse secondary follicles. **A**. Schematic representation of microinjection and *in vitro* oocyte growth culture. The secondary follicles were cultured for 1 d, subsequently microinjected mRNA and/or siRNA, and cultured for another 15 d to produce transfected fully grown oocytes. **B**. Cytoplasmic injection to the growing oocyte in a secondary follicle. Upper, before microinjection; lower, microinjecting. **C**. mCherry expression in the secondary follicles at 2 d post microinjection. Scale bars indicate 50 μm.

After mRNA microinjection, the 2-step IVG culture was performed using a modified version of the method developed previously [18]. Fully grown oocytes were obtained after 5 days of the first-step culture followed by 10 days of the second-step culture.

mCherry fluorescence was maintained throughout the IVG culture period but declined sharply immediately after fertilization (**Fig. 2A–B**). Notably, mCherry fluorescence was detectable from the oocyte growth stage through the 2-cell stage; however, no statistically significant difference was observed between the no injection and mCherry mRNA injection groups at the blastocyst stage (P = 0.999). In mouse embryos, fertilization triggers extensive degradation of maternal factors together with large-scale reorganization of proteins and mRNAs through autophagy and zygotic genome activation [19–21]. The rapid disappearance of mCherry fluorescence after fertilization therefore suggests that the exogenous mCherry protein or its transcripts may be degraded in a manner similar to endogenous maternal factors. Because the microinjection procedure physically generates a minute perforation in the follicle by microinjection, growing oocytes or surrounding granulosa cells occasionally leaked out of the follicle through the injection site, likely due to intrafollicular pressure (**Fig. 2C**). Despite exhibiting a certain degree of cellular damage, microinjected follicles continued to grow and displayed early embryonic developmental competence comparable to that of the no injection group following fertilization (Table 1). These findings suggest that the cytoplasmic microinjection of mRNA into growing oocytes in the secondary follicles enables overexpression of exogenous genes while largely preserving the developmental competence of oocytes. Nevertheless, because the current data are preliminary, further studies will be required to rigorously assess the extent of cellular damage associated with the microinjection procedure.

**Figure 2.**
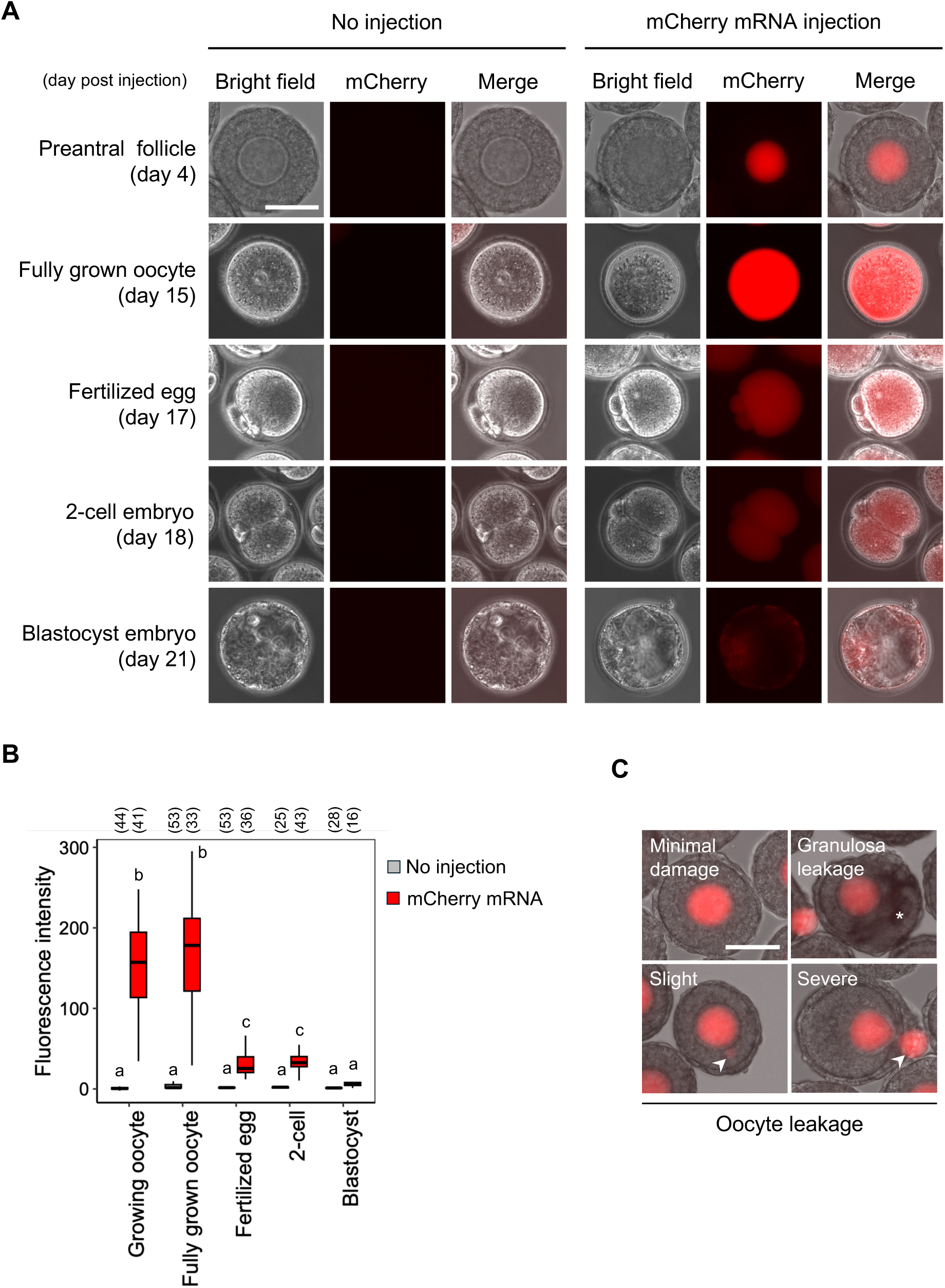
Experimental design for microinjection and *in vitro* growth of mouse secondary follicles. **A**. mCherry expression of microinjected oocytes and follicles during oocyte growth and preimplantation development. **B**. Relative intensity of mCherry fluorescence in oocytes and embryos. ^a,b,c^Groups with different letters indicate significant differences (Tukey’s test, *P* < 0.01). **C**. Microinjection-induced damage to growing oocytes and follicles. Asterisk, leaked granulosa cells; arrowhead, leaked oocytes. Scale bars indicate 50 μm.

**Table 1.** *In vitro* oocyte growth and post-fertilization development of microinjected oocytes.

|  | No. of<br>injected<br>follicle | No. of<br>enlarged<br>follicle<br>(day 16) | No. of<br>expanded<br>COC<br>(day 17) | No. of<br>fertilized<br>oocyte<br>(day 18) | No. of<br>2-cell<br>embryo<br>(day 19) | No. of<br>blastocyst<br>embryo<br>(day 22) |
| --- | --- | --- | --- | --- | --- | --- |
| No injection | 120 | 77 (64.2%) | 65 (84.4%) | 46 (70.8%) | 43 (93.5%) | 30 (69.8%) |
| Injection | 83 | 60 (72.3%) | 46 (76.7%) | 26 (56.6%) | 24 (92.3%) | 15 (62.5%) |

Next, we suppressed DNA methyltransferase 3-like (*Dnmt3l*) gene to evaluate whether the KD efficiency was sufficient to silence a highly expressed endogenous gene. *Dnmt3l* is expressed in oocytes during oocyte growth phase as well as in several embryonic stages and interacts with other DNA methyltransferases to facilitate *de novo* DNA methylation [22, 23]. Maternal *Dnmt3l* KO oocytes normally grow to the fully grown stage and can be fertilized; however, the resulting embryos exhibit lethality during mid-gestation accompanied by global loss of DNA methylation across the genomes, including differentially methylated regions of maternal imprinted genes. Cultured oocytes microinjected with siRNAs targeting *Dnmt3l* together with mCherry mRNA grew normally to the fully grown stage and exhibited mCherry fluorescence throughout the oocyte growth period (**Fig. 3A–B**). Only mCherry-positive oocytes were collected under a fluorescence microscope and analyzed as *Dnmt3l* KD oocytes. A key advantage of this method is that co-injecting fluorescent reporter mRNA with siRNA allows for the selective exclusion of oocytes that failed to be injected properly due to technical errors or insufficient volume of the mixture. This screening process ensures that only successfully and adequately microinjected oocytes are used for subsequent analysis, thereby enhancing the reliability of the knockdown experiments. Most importantly, endogenous *Dnmt3l* mRNA expression was markedly reduced in *Dnmt3l* KD oocytes (**Fig. 3C**; siNegative = 100% vs. si*Dnmt3l* = 5.5%). These results indicate that this method efficiently suppressed endogenous gene expressions throughout the oocyte growth phase without detectable growth impairment.

**Figure 3.**
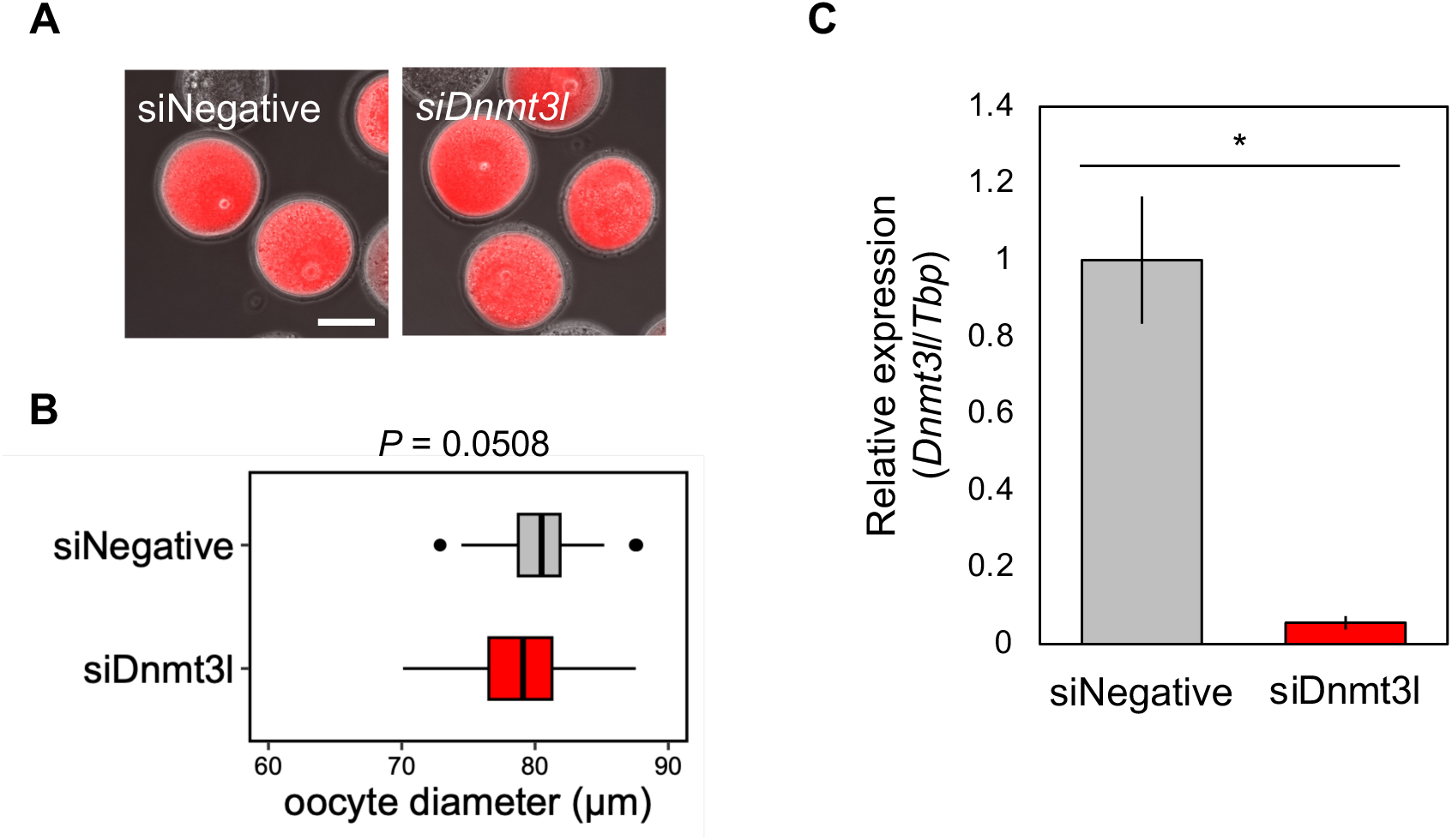
Knock-down effect in microinjected and *in vitro*-grown oocytes. **A.** mCherry expression in mCherry mRNA and siRNA for *Dnmt3l*-microinjected fully grown oocytes. Scale bar indicates 50 μm. **B**. Oocyte diameter of siRNA for *Dnmt3l*-delivered fully grown oocyte at 16 d of culture (siNegative, N = 45; si*Dnmt3l*, N = 35). **C**. Relative expression of *Dnmt3l* for siRNA-microinjected fully grown oocytes (each N = 4).

In the CKO strategy using mice using mice expressing Cre recombinase specifically in female germ cells, the target gene is irreversibly disrupted during oogenesis, meaning its function cannot be restored after fertilization. In contrast, our strategy is designed to inhibit gene function specifically during the oocyte growth phase. Since the introduced siRNAs are gradually diluted and/or degraded through subsequent embryo development and cell proliferation, the target gene expression is expected to recover to normal levels from the intact genome.

The present study introduces another approach that enable rapid *in vitro* evaluation of maternal effect gene function without interfering with early embryonic development, providing a significant advantage for functional analysis of maternal factor within a 16-day culture period. Furthermore, this system facilitates the analysis of combinatorial gene functions through the introduction of multiple siRNAs into growing oocytes, making it a versatile platform for studying not only oogenesis but also fertilization as well *in vitro*.

## Materials and methods

### Ethics

All animal experiments were conducted under approval of the Gunma University Institutional Animal Care and Use Committee (approval number 21-025), according to the Guidelines for Proper Conduct of Animal Experiments by the Science Council of Japan.

### Mice

We purchased C57BL/6N and DBA/2 from and SLC Japan (Hamamatsu, Japan) and maintained them in the Bioresource Center at Gunma University. B6D2F1 mice were obtained by mating C57BL/6N female and DBA/2 male mice. B6D2F1 female mice at P10 were sacrificed for secondary follicle collection. Mature B6D2F1 male mice were also sacrificed for spermatozoa collection.

### Microinjection to secondary follicles

Microinjection experiment was conducted as a previous report with minor modifications [12]. Secondary follicles were isolated from ovaries of B6D2F1 female mice at P10, using 28 G needle in L-medium (Sigma-Aldrich, St. Louis, MO, USA) containing 0.4% (w/v) polyvinylpyrrolidone (PVP; 360 kDa, P5288, Sigma-Aldrich). Five secondary follicles were cultured in a 5 μl droplet of α-MEM, GlutaMAX Supplement (Gibco, Thermo Fisher Scientific, Waltham, MA, USA) containing 10% (v/v) fetal bovine serum (FBS; Thermo Fisher Scientific) on a 35-mm petri dish (1008; Corning Inc., Corning, NY, USA) to remove ovarian interstitial cells at 37°C, 5% CO2 in air overnight. We injected each 25 μM siRNA for *Dnmt3l* (si*Dnmt3l*-1, SASI_Mm02_002990031; si*Dnmt3l*-2, SASI_Mm02_002990033, Merck, Darmstadt, Germany) and 500 ng/μl mCherry mRNA into the ooplasm using FemtoJet 4i (Eppendorf, Hamburg, Germany). mCherry mRNA was synthesized using mMESSAGE mMACHINE T7 Transcription Kit (Invitrogen, Thermo Fisher Scientific) and Poly(A) Tailing Kit (Invitrogen) as template mCherry coding sequences cloned into pGEM-T Easy Vector (Promega, Madison, WI, USA). Injection protocol was below: injection pressure, 1000 hPa; injection time, 0.5 sec; compensation pressure, 100 hPa. Once injected into a secondary follicle, we rinsed the capillary by clean mode after each microinjection. Injection capillaries (Borosilicate Glass with Filament, BF100-78-10, Sutter Instrument, Novato, CA, USA) were pulled using a PC-100 Puller (Narishige, Tokyo, Japan) with two-stage condition (stage 1 and 2: 65 heater level 4 x weight), and the tip of pulled injection capillaries was carefully fractured to create a small opening using MF2 Micro Forge (Narishige). Holding pipettes (Borosilicate Glass, B100-75-10, Sutter Instrument) were pulled using a PC-100 with one-stage condition (70 heater level, 4 x weight). The tip of pulled capillaries was cut and fire-polished using MF2 Micro Forge (Outer thickness = 100–110 μm, inner thickness = 35–45 μm).

### *In vitro* oocyte growth

We performed two-step IVG as previous reports with minor modifications [16–18]. Injected follicles were cultured in α-MEM (Gibco) supplemented 2% (w/v) PVP, 5% (v/v) FBS, 0.1 IU/ml human recombinant follicle stimulating hormone (FSH; Gonalef; Merck), and 0.1% (v/v) penicillin-streptomycin solution (P4458; Sigma-Aldrich) for another 15 days to grow to the antral stage. First, follicles were cultured in on a Millicell cell culture insert (PICM0RG50; Merck) for 5 days in a 35-mm petri dish under 37°C, 5% CO2, and 20% O2 condition. The medium was added at 1 ml inside and 2 ml outside the Millicell insert, respectively. During the semi-floating culture in the Millicell membrane, half of the culture medium was replaced with fresh medium at the day 3 of culture. On the day 5 of culture, microinjected follicles were treated with 0.1% (w/v) collagenase type I (Worthington Biochemicals, Lakewood, NJ, USA) in L-15 medium containing 30% (v/v) FBS for 30 min at 37°C, 5% CO2, and 20% O2 condition to remove the theca cell layers and washed in L-15 medium containing 0.4% PVP. Up to 70 theca-free follicles were cultured on collagen-coated Transwell membrane (3414; Corning Inc.) in a 6-well plate under 37°C, 5% CO2, and 7% O2 condition for additional 10 days (total days). The medium was added at 1.5 ml inside and 2 ml outside the Transwell insert, respectively. During the adherent culture in the Transwell membrane, half of the culture medium was replaced with fresh medium every other day.

### *In vitro* maturation and *in vitro* fertilization

COCs from cultured follicles at IVG14d were cultured for 17 h in α-MEM containing 5% (v/v) FBS, 0.1 IU/ml FSH, 1.2 IU human chorionic gonadotropin (Gonatropin; ASKA Pharmaceutical, Tokyo, Japan), and 4 ng/ml epidermal growth factor (Gibco) under 37°C, 5% CO2, and 20% O2 condition [24]. The cumulus-expanded COCs were fertilized with capacitated sperm in TYH medium (LSI Medience, Tokyo, Japan) for 6 h under 37°C, 5% CO2, and 20% O2 condition [9]. Fertilized oocytes with two pronuclei were cultured in EmbryoMax KSOM Mouse Embryo Media (Merck) for 5 days under 37°C, 5% CO2, and 20% O2 condition.

### Fluorescence measurement

During cultivation, mCherry fluorescence of injected oocytes and embryos was observed with a BZ-X700 fluorescent microscopy (Keyence, Osaka, Japan). mCherry signal intensities relative to the growing oocytes of no injection group was measured using ImageJ software [25].

### Gene expression analysis

Total RNA was extracted from oocytes and embryos using an RNeasy Micro Kit (Qiagen, Venlo, Netherland) according to the manufacturer’s instructions, and reverse-transcribed into cDNA using PrimeScript RT Reagent Kit with gDNA Eraser (Takara Bio, Kusatsu, Japan) with Oligo(dT) 12–18 primer (Thermo Fisher Scientific). We performed quantitative RT-PCR using THUNDERBIRD SYBR qPCR Mix (Toyobo, Osaka, Japan) on a StepOnePlus Real-Time PCR System (Applied Biosystems, Thermo Fisher Scientific) to determined relative expression levels of *Dnmt3l* and *Tbp* (as internal control). The sequences of primer pairs for Gene expression analysis were as follows: *Dnmt3l*, 5′-CTAGCCCTGATGCTGACAGT-3′ (fwd) and 5′-AGGCTCTTTGCAGTCTTCCA-3′ (rev); *Tbp*, 5′-ATCCCAAGCGATTTGC-3′ (fwd) and 5′-GCTCCCCACCATGTTC-3′ (rev) [17]. Relative expression level was calculated by ddCt methods [26]. The significant difference was assessed by t-test (*P* < 0.01).

## Acknowledgements

We are grateful to the staffs at the Bioresource Center of Gunma University Graduate School of Medicine to their contributions to animal cares. This work was supported by JSPS KAKENHI (grant number 24K09199 to KS), by the Basic Science Research Project of the Sumitomo Foundation (grant number 2402483 to KS). This work was also the result of using research equipment shared in the MEXT Project for promoting public utilization of advanced research infrastructure (Program for supporting the introduction of the new sharing system, JPMXS0420600123).

## Author contribution

**A.** K. S., Y. S. and M. M. designed experimental procedures and performed all experiments. K. S., Y. S., A. J-O. and T. M. prepared and reviewed the manuscript.

